# Individual differences in response precision correlate with adaptation bias

**DOI:** 10.1101/285973

**Authors:** Marcelo G. Mattar, Marie V. Carter, Marc S. Zebrowitz, Sharon L. Thompson-Schill, Geoffrey K. Aguirre

**Affiliations:** Department of Psychology, University of Pennsylvania, Philadelphia, PA 19104, USA; Department of Neurology, University of Pennsylvania, Philadelphia, PA 19104, USA

**Author notes:** Present address: Princeton Neuroscience Institute, Princeton University, Princeton, NJ 08544, USA.

**Keywords:** visual adaptation, individual differences, face perception, color perception, Bayesian inference, Amazon Mechanical Turk

## Abstract

The internal representation of stimuli is imperfect and subject to bias. Noise introduced at initial encoding and during maintenance degrades the precision of representation. Stimulus estimation is also biased away from recently encountered stimuli, a phenomenon known as adaptation. Within a Bayesian framework, greater biases are predicted to result from poor precision. We tested for this effect in individual difference measures. 202 subjects contributed data through an on-line experiment. During separate face and color blocks, subjects performed three different tasks: an immediate stimulus-match, a 5 seconds delayed match, and 5 seconds of adaptation followed by a delayed match. The stimulus spaces were circular and subjects entered their responses on a color/face wheel. Bias and precision of responses were extracted while accounting for the probability of random guesses. We found that the adaptation manipulation induced the expected bias in responses, and the magnitude of this bias varied reliably and substantially between subjects. Across subjects, there was a negative correlation between mean precision and bias. This relationship was replicated in a new experiment with 192 subjects. This result is consistent with a Bayesian observer model, in which the precision of perceptual representation influences the magnitude of perceptual bias.

## Introduction

Perception is our “best guess” as to what is in the world (von Helmholtz, 1867). When sensory information is imperfect or incomplete, this best guess relies more heavily on expectations and prior beliefs (Knill and Richards, 1996). Prior knowledge influences perception continuously and automatically, and may occasionally lead to perceptual illusions (Bar, 2004; Summerfield and Egner, 2009; Lafer-Sousa et al., 2015). To achieve statistically optimal inference, sensory evidence and prior beliefs are quantitatively combined using Bayes’ rule (Bayes and Price, 1763).

While many aspects of human perception are found to be consistent with Bayesian observer models (Knill and Richards, 1996; Stocker and Simoncelli, 2006b), it has been more difficult to apply this framework to the ubiquitous phenomenon of sensory adaptation. Recent sensory history tends to bias perception “away” from observed stimuli (Levinson and Sekuler, 1976; Clifford, 2002, although see Fischer and Whitney, 2014 and Gibson and Radner, 1937), which seems at at odds with a Bayesian account of perception that is biased toward prior beliefs. Recent theoretical work attempts to explain perceptual adaptation as arising either from an asymmetric likelihood (Wei and Stocker, 2015) or a modification of the prior (Chopin and Mamassian, 2012). In either case, larger perceptual biases should be observed in the setting of greater imprecision.

Here, we test this key prediction of Bayesian inference. Using a web-based visual adaptation experiment with color and face stimuli, we investigated the relationship across individuals between variability in response precision and variability in adaptation biases. We found that precision and bias are relatively stable measures of an individual, and that biases for color stimuli are correlated with biases for face stimuli. We then investigated the relationship between bias and precision across individuals and found that biases for both materials are larger when precision is lower. These results conform with predictions of Bayesian observer models whereby perception is biased away from recently observed stimuli when sensory information is uncertain.

## Methods

### Participants

1002 people were recruited through Amazon Mechanical Turk, 530 for the main experiment (Table 1) and 472 for the replication experiment (Tables 3 and 4). This research was reviewed and deemed exempt from oversight by the University of Pennsylvania Institutional Review Board, and therefore informed consent was not collected. Information on the home page of the web-based experiment indicated the research nature of the project. No information that could identify participants was collected. All subjects received a fixed minimum compensation of $0.25 for their participation in addition to a performance-based bonus of up to $12.00. The full experiment took approximately 1 hour and 15 minutes to complete, and subjects that reached the end received an average bonus of $7.68 ($0.51 SD).

**Table 1:** Subject enrollment and exclusion in the main experiment. The total in the cells may not match the total number of subjects due to missing responses from some subjects.

| Completed: | Registration | Screening | Experiment |
| --- | --- | --- | --- |
| Number of subjects | 530 | 422 | 202 |
| Age: $M \pm SD$ | $36 \pm 12$ | $36 \pm 11$ | $35 \pm 11$ |
| Male/Female | 252/277 | 191/231 | 92/110 |
| Left/Right-handed | 18/508 | 14/405 | 3/197 |

**Table 2:**
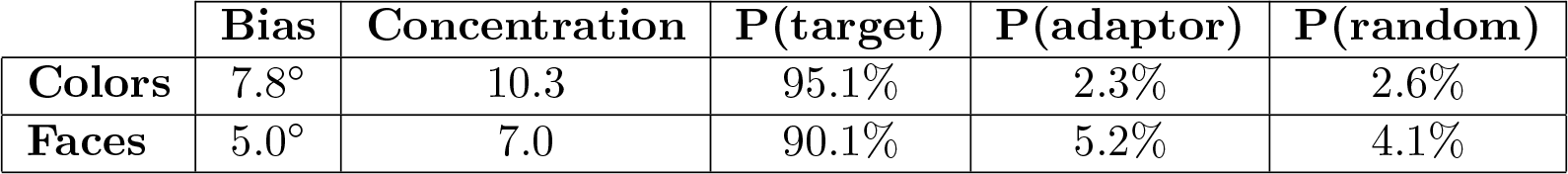
Parameters from group fit of the adaptation data in the main experiment.

|  | Bias | Concentration | P(target) | P(adaptor) | P(random) |
| --- | --- | --- | --- | --- | --- |
| **Colors** | 7.8° | 10.3 | 95.1% | 2.3% | 2.6% |
| **Faces** | 5.0° | 7.0 | 90.1% | 5.2% | 4.1% |

**Table 3:** Subject enrollment and exclusion in the replication experiment with colors. The total in the cells may not match the total number of subjects due to missing responses from some subjects.

| Completed: | Registration | Screening | Experiment |
| --- | --- | --- | --- |
| Number of subjects | 191 | 149 | 98 |
| Age: $M \pm SD$ | $37 \pm 12$ | $37 \pm 12$ | $39 \pm 12$ |
| Male/Female | 110/81 | 87/62 | 60/38 |
| Left/Right-handed | 8/176 | 4/139 | 0/95 |

**Table 4:** Subject enrollment and exclusion in the replication experiment with faces. The total in the cells may not match the total number of subjects due to missing responses from some subjects.

| Completed: | Registration | Screening | Experiment |
| --- | --- | --- | --- |
| Number of subjects | 281 | 234 | 94 |
| Age: $M \pm SD$ | $36 \pm 12$ | $37 \pm 12$ | $37 \pm 12$ |
| Male/Female | 164/117 | 134/100 | 48/46 |
| Left/Right-handed | 19/262 | 13/221 | 3/91 |

### Stimuli and Materials

The experiment was programmed in JavaScript and hosted on a custom-built website that subjects accessed using their own computers (https://cfn.upenn.edu/iadapt). Stimuli consisted of synthetic faces generated with FaceGen Main SDK (Inversions, 2012) and colored squares.

The face set used in the main experiment varied in age and gender. Two base stimuli were generated by varying the gender of an identity-neutral face from male to female, and another two base stimuli were generated by varying the age of an identity-neutral face from 15 to 65 years old. Based on these four stimuli, a set of 360 faces were generated in a circular space by varying both dimensions, with main axes corresponding to age and gender.

The color set varied in hue with no nominal variation in lightness and saturation. A set of 360 color values were generated in HSL space with L* held fixed at 25 and saturation equal to 7. This saturation value was determined in pilot experiments as producing stimuli whose response variance matched approximately that of face stimuli. The set of HSL value were then converted to sRGB space.

For the replication experiment, new sets of face and color stimuli were generated. The face stimuli were again generated based on four stimuli that varied by apparent age (15-65 years old) and gender (male-female). To produce more distinctive faces than the previous set, the four base stimuli also varied in internal facial features. This was realized by displacing each base face stimulus by along a random vector direction within the face stimulus space, orthogonal to both age and gender. The color stimuli were again generated in HSL space with L* held fixed at 25, but now with saturation equal to 20 (the maximum value that produced sRGB values within the 0-255 range displayable in regular monitors), producing more distinctive colors than in the main experiment.

### Experimental procedure

Subjects recruited through Mechanical Turk were redirected to the experiment website, where they entered their responses using mouse and keyboard input. Upon completion, subjects received an anonymous code that they provided to Mechanical Turk for payment. Each experimental session started with a basic description of the procedures, followed by a demographics questionnaire. Subjects then completed a screen calibration procedure and a color perception test followed by the main experimental blocks.

In the screen calibration procedure, subjects were presented with a set of discrete color gradients organized in rows, each ranging from black to a distinct, saturated color value. They were then asked to adjust the screen settings and/or the angle of their laptop screen to allow them to simultaneously distinguish between neighboring colors on both ends of the each gradient. In the color perception test, subjects completed eight trials of the Ishihara test, a test for congenital color deficiencies. Subjects proceeded to the main experimental blocks only if at least seven responses were correct. This on-line test was not capable of detecting subtle color vision deficits, but did ensure that the subject and their computer could support a baseline level of color discrimination ability.

Subjects then completed a battery of tests designed to estimate representation precision, precision after a delay, and precision and bias after adaptation. Data were collected over 2-6 blocks of each of the following experiments: (i) *stimulus-match*; (ii) *delayed-match*; (iii) *adaptation* (Fig. 1a). Results from the delayed-match task are not further considered here. Each block consisted of 15-30 trials (each lasting between 2 and 5 minutes) in which subjects were instructed to report the value of a target stimulus — a color or a face — by clicking and moving the mouse cursor around a stimulus wheel (Fig. 1b), allowing a fine adjustment of their responses. Prior to performing each type of experiment for the first time, subjects were presented with detailed instructions, a mini-quiz containing three questions with three alternatives each about the instructions, and five practice trials. If any answer to the quiz was incorrect, subjects were repeatedly presented with the instructions and the quiz, until all three answers were simultaneously correct. Similarly, subjects repeated the practice experiment as many times as necessary until all five responses were within 5° of the target. Together, these approaches ensured comprehension of the experiment instructions, and that subjects were able to adequately perform the experiment on their computer.

**Figure 1:**
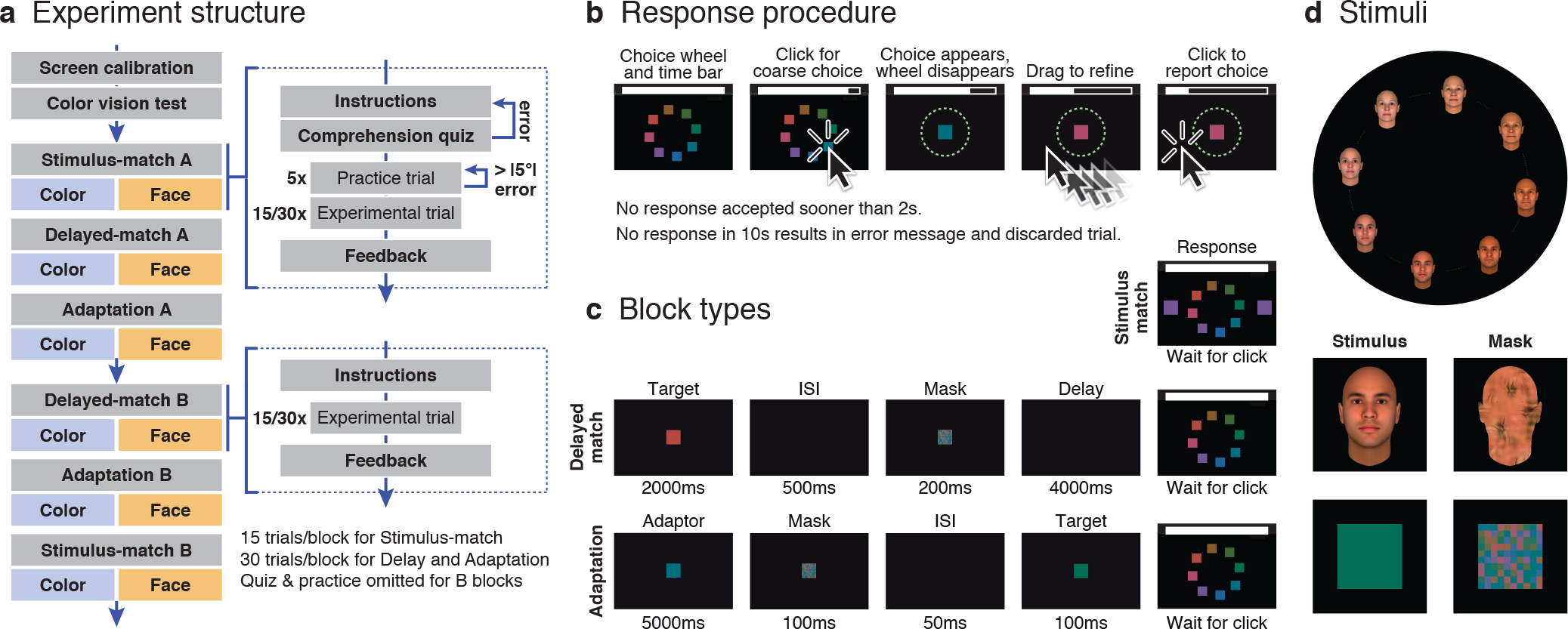
Experiment setup and methods. *(a)* Experiment structure. Each subject completed two blocks of each experiment, for both color and face stimuli. Prior to each experiment, subjects were presented with instructions and a comprehension quiz; the first time they performed each block of a given experiment, they also completed 5 practice trials. Each block of the stimulus-match experiment had 15 trials. Each block of the delayed-match and adaptation experiments had 30 trials. *(b)* Response procedure. Subjects were first presented with 8 equally spaced representative stimuli around the wheel. To enter their responses, subjects first performed a coarse selection by clicking on the region of the wheel that approximately matched the target. They then adjusted their selection more precisely by dragging the pointer around the wheel and clicking a second time to confirm their responses. *(c)* Block types. Top: On each trial of the stimulus-match experiment, subjects were instructed to match to a stimulus presented on both sides of the screen; the target stimulus remained on the response screen while the subject made the match. Middle: On each trial of the delayed-match experiment, subjects were instructed to match to a target stimulus following a 4 second interval. The data from this task are not further considered in this report. Bottom: On each trial of the adaptation experiment, subjects were instructed to match to a target stimulus that was presented after a 5 second adaptation period and a mask. No interval separated adaptor and mask, and an interval of 50 ms separated mask and target (not shown). *(d)* Stimuli. A circular space with 360 stimuli was used for both color and face stimuli. Color stimuli were generated to vary in hue but not in saturation or luminance (HSL space). Face stimuli varied in age and gender, each forming one axis of the space. Color masks were a checkerboard composed of various stimuli randomly sampled from the color space. The eight stimuli that are snown in the figure and in the response screen are equally spaced examples from the entire set of 360, and were selected at random on each trial. Face masks were created using the steerable pyramids method to match various low-level visual properties (Portilla and Simoncelli, 2000).

Experimental blocks in the main experiment were completed in the following order: (1) color stimulus- match; (2) face stimulus-match; (3) color delayed-match; (4) face delayed-match; (5) color adaptation; (6) face adaptation; (7) color delayed-match; (8) face delayed-match; (9) color adaptation; (10) face adaptation; (11) color stimulus-match; (12) face stimulus-match (Fig. 1a). In the replication experiment, blocks (only one stimulus class) were completed in the following order: (1) stimulus-match; (2) delayed-match; (3-8) adaptation; (9) delayed-match; (10) stimulus-match (Fig. 6a).

We calculated subject accuracy on every trial by linearly mapping from errors (0° – 90°) to accuracy (100% - 0%) and, at the end of each block, we calculated the average accuracy for that block (100% being perfect performance). The compensation accumulated by subjects increased at the end of each experimental block by an amount proportional to the average accuracy. Subjects were presented with their average accuracy in the finished block, the corresponding dollar amount accumulated, and the total compensation accumulated in the experimental session up to that point. If the accuracy on any block ended up below 20%, the experimental session was terminated and subjects were directed back to Amazon Mechanical Turk to receive their payment. Only subjects who maintained accuracy above 20% in all blocks were able to reach the end of the experiment; those subjects received twice the regular compensation. Out of the 1002 subjects recruited, 197 were excluded for either abandoning or not passing the color perception test, and 411 for not maintaining accuracy above 20% throughout the entire session. Only data from the remaining 394 subjects were included in the analyses described in this paper (Tables 1, 3, 4).

### Stimulus-match experiment

Each block consisted of 15 trials, with target values sampled uniformly (24° spacing) from the circular space; each target was presented once per block. On each trial, subjects were presented with a target stimulus on both the left and right sides of the black background screen, along with a stimulus wheel containing eight, equally spaced thumbnails with representative stimuli from the circular space (Fig. 1*c*, *top*). Because we could not control the properties of the display used to present the stimulus to each subject, we could not ensure that the perceptual similarity between equally spaced stimuli was uniform around the wheel. Further, the perception of the stimuli may be subject to long-term priors over the stimulus space. However, by sampling uniformly from the wheel and providing test stimuli in both the clockwise and counter-clockwise direction, we expect either of these forms of stimulus bias to be ameliorated on average. Moreover, the specific thumbnails, their position, and the mapping of stimulus value to screen position, varied randomly on each trial (i.e., there was no systematic relationship between stimulus values and screen location). Subjects were instructed to click once with the cursor positioned on the region of the screen corresponding to the target location. The selected stimulus was then presented in the center of the screen, and subjects were allowed to fine-tune their response by moving the cursor around the stimulus wheel before confirming their selection with a second mouse click (Fig. 1b).

During the fine-tuning phase, the stimulus presented in the center of the screen varied (in steps of 1°), to allow subjects to precisely match their responses to the target stimulus. The second (confirmation) click was registered only if it occurred within 2-10 seconds after the trial onset. If no response was entered for 10 seconds, a dialog box was displayed warning the subject to pay attention and click the “OK” button to continue. On these trials, an accuracy of 0% was registered (for the purpose of calculating the average block accuracy), though they were not included in the main analyses. Similarly, if responses were more than 90° away from the target, a modal dialog box was displayed warning the subject to pay attention. These measures ensured that subjects maintained continuous attention throughout the entire block and slowed down subjects who attempted to rush through the experiment without care.

### Delayed-match experiment

Each block consisted of 30 trials, with target values sampled uniformly (12° spacing) from the circular space — i.e., each target was presented once per block. On each trial, subjects were presented with a target stimulus in the center of the screen for 2000 ms, followed by a mask stimulus (ISI: 500 ms) at the same location for 200 ms, followed by a 4000 ms interval of a blank screen during which no response was allowed (Fig. 1c, Middle). After this interval, subjects entered their responses using the same procedure described previously (Fig. 1b). The same measures described previously were used to ensure that subjects maintained continuous attention throughout the entire block.

Color masks were checkerboards composed of various colors randomly sampled from within the stimulus set. Face masks were created by synthesizing textures based upon the original face stimuli using the steerable pyramid approach (Portilla and Simoncelli, 2000; Fig. 1d). The resulting textures were placed within the outlines of the face stimuli, resulting in a mask that had similar low level feature properties to the faces (e.g., spatial frequency, line curvature) but was not recognizable as a face.

### Adaptation experiment

Each block consisted of 30 trials, with target values sampled uniformly (12° spacing) from the circular space. Each target was presented once per block. On each trial, subjects were presented with an adaptor stimulus in the center of the screen for 5000 ms, drawn from a position in the stimulus space ±45° from the target (adaptors at +45° and −45° were intermixed within a block). The adaptor was immediately followed by a mask stimulus at the same location for 100 ms, by target stimulus for 200 ms (ISI: 50 ms), and by a 100 ms interval of a blank screen during which no response was allowed (Fig. 1c, Bottom). After this interval, subjects entered their responses using the same procedure described previously (Fig. 1b). Subjects received a warning if their responses were within 10° of the adaptor stimulus position. In these cases, a modal dialog box was presented indicating that subjects should ignore the adaptor and report the value of the target.

### Data analysis

We used methods for circular data (Fisher, 1995). We calculated the error on each trial as the angular deviation on the stimulus wheel between the target value and the response entered. We then used maximum likelihood estimation to fit the distribution of error values in the circular space. In both stimulus-match and delayed-match experiments, the distribution of error values was decomposed into two parameters that represent a mixture of a uniform distribution (corresponding to random responses) and a von Mises distribution — the circular analog of the Gaussian distribution on a line — centered on the target value. The parameters fit by this procedure correspond to the probability of guesses, which is proportional to the height of the uniform distribution, and the precision of responses, which is the inverse of the standard deviation of the best-fitting Gaussian distribution (which is in turn related to the concentration parameter of the fitted von Mises distribution; Fig. 2a).

**Figure 2:**
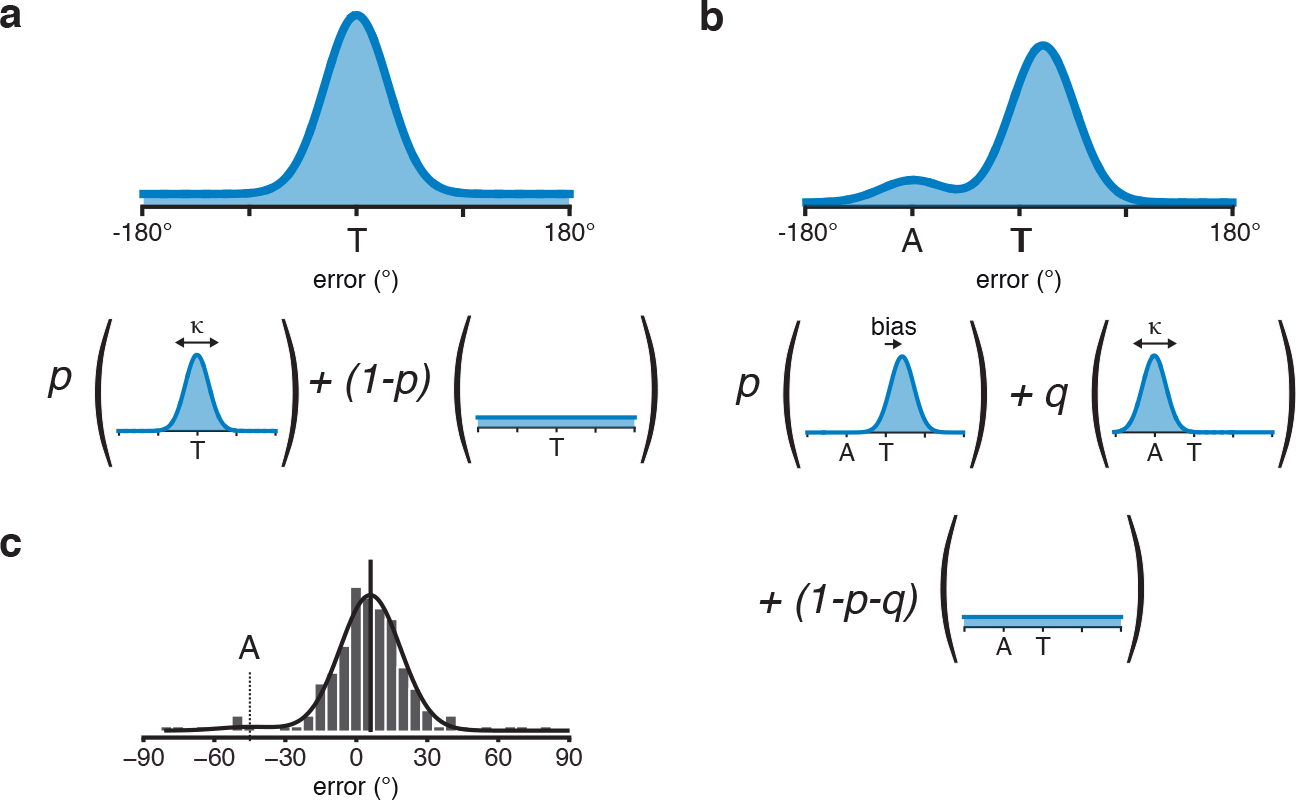
Mixture model fitting approach. The distribution of errors was modeled as a superposition of probability distributions to account for different types of responses. *(a)* In the stimulus-match and delayed-match experiments, responses could be concentrated near the target value (von Mises distribution centered at the target value) or fall randomly in any position of the space with equal probability (von Mises distribution with concentration parameter equal to zero, equivalent to a uniform distribution). Two parameters were estimated: the concentration parameter of the responses near the target, and the probability of random responses. *(b)* In the adaptation experiment, responses could be concentrated near the target value (von Mises distribution displaced from the target by a fixed amount), concentrated near the adaptor value (a stimulus that the subjects were instructed to ignore, modeled as a von Mises distribution centered at the adaptor value) or fall randomly in any position of the space with equal probability (von Mises distribution with concentration parameter equal to zero). Four parameters were estimated: the magnitude of the displacement of responses near the target (bias), the concentration parameter of the responses near the target, the probability of responses near the adaptor, and the probability of random responses. *(c)* Example of a good fit for a subject in the adaptation experiment. Data are collapsed across both blocks of the adaptation experiment, and indicates the existence of a positive (repulsive) bias in relation to the adaptor.

In the adaptation experiment, the distribution of error values was decomposed into four parameters that represent a mixture of a uniform distribution (corresponding to random responses), a von Mises distribution centered on the adaptor value (corresponding to responses where the subject mistakenly attempts to report the value of the adaptor due to not seeing or not paying attention to the target), and a von Mises distribution with equal concentration parameter centered *near* the target value. The parameters fit by this procedure correspond to the probability of guesses, the precision of responses (the inverse of the standard deviation of the best-fitting Gaussian distribution), and the bias, which is the mean of the von Mises distribution centered near the target value (Fig. 2b).

## Results

We investigated the relationship between variation in precision and adaptation bias across 530 individuals. Each subject performed a series of psychophysics experiments on their personal computers and received compensation for their time (Fig. 1a). On each experimental trial, subjects were instructed to report the value of a target stimulus — a color or a face — by clicking and dragging the mouse pointer around a stimulus wheel, allowing a fine adjustment of their responses (Fig. 1b). Wheels were comprised of 360 distinct stimuli varying in hue (colors) or in age and gender (faces; Fig. 1d).

Subjects completed three types of experiments for each stimulus class. In the *stimulus-match* experiment (15 trials), designed to obtain a baseline response precision for each subject, subjects were instructed to select a value on the wheel matching the target stimulus (Fig. 1c, top). In the *delayed-match* experiment (30 trials), designed to estimate subject’s working memory precision, a target stimulus was presented on the center of the screen for 2.0 s, followed by a 4.0 s delay, after which subjects were to select a value on the wheel matching the target stimulus (Fig. 1c, middle). In the *adaptation* experiment (30 trials), designed to estimate the magnitude of adaptation biases, an adapting stimulus was presented in the center of the screen for 5 seconds followed by brief mask and a target stimulus ±45° away from the adaptor for 200ms, after which subjects were to select a value on the wheel matching the target stimulus (Fig. 1c, bottom). Subjects performed two separate blocks of each experiment in a session, allowing for tests of measure reliability, and the trials within each experiment were sampled uniformly and in random order from the circular space (Fig. 1d). Throughout the entire session, subjects were only allowed to proceed to the next block of the experiment if their accuracy remained above a minimum threshold (see Methods). A total of 328 subjects were either excluded or abandoned the experiment (108 at the color vision test and 220 during the main experimental trials), leaving 202 subjects for the main analyses described in this chapter (Table 1).

We calculated the error on each trial as the difference between the target value and the response entered (*ε* = *θ*_*target*_ − *θ*_*response*_), and we fit the distribution of error values for each subject using a superposition of distributions defined over a circular support (−180° < *θ* < +180°) (Zhang and Luck, 2008; Bays et al., 2009). This procedure simultaneously estimates the precision of the error distribution (i.e., the inverse of the standard deviation of the best-fitting Gaussian distribution; Jammalamadaka and Sengupta, 2001) and the probability of random guesses (Fig. 2a).

In the adaptation experiment, biases expected to result from adaptors located at +45° and −45° away from the target should have equal magnitude and opposite sign. Thus, we combined the distribution of error values from trials where the adaptor was located at −45° from the target (*ε*) with the distribution of reflected error values (−*ε*) from trials where the adaptor was located at +45° from the target. In these experiments, the fitting procedure also estimates two additional parameters: the mean of the error distribution (i.e., the bias induced by the paradigm) and the probability that the selected response matches the adaptor stimulus and not the target stimulus (Fig. 2b,c).

### Adaptation produces a repulsive bias

To confirm the effectiveness of our adaptation paradigm in inducing repulsive biases, we first fit the adaptation data from all subjects combined (12,120 trials per stimulus class) with a mixture of distributions as described. We observed a positive (repulsive) bias of 7.8° and 5.0° for color and face stimuli, respectively. We also estimated that subjects responded randomly in 2.6% and 4.1% of the trials, and that their responses matched the adaptor stimulus in 2.3% and 5.2% of the trials, with the remaining 95.1% and 90.8% of responses concentrated around the target value (Fig. 3). These results confirm that the paradigm induced the typical *repulsive* after-effects, in which subject responses to target stimuli tend to be biased *away* from the preceding, adapting stimulus (Table 2).

**Figure 3:**
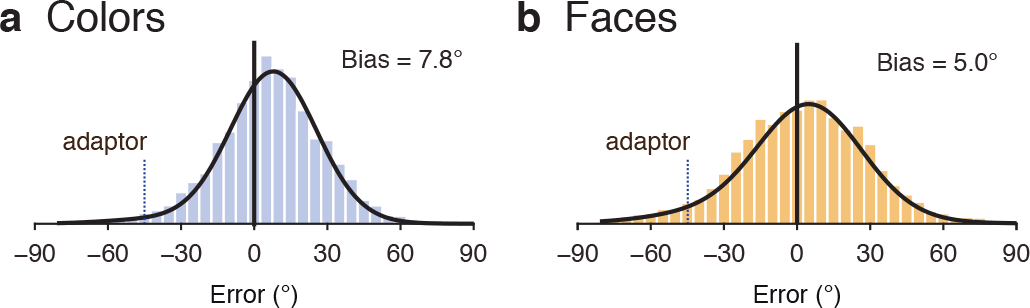
Estimation of adaptation biases at the population level. *(a)* Group adaptation results for all color trials. The bars provide the histogram of error (5° bins) in responses across all trials and all subjects in the adaptation experiment that used color stimuli. If subjects reported the color of the target stimulus perfectly, all trials would have zero error. Each trial in the adaptation experiment featured a five second adaptor stimulus, which in this plot has a relative location of −45° and is indicated with the blue dotted line. The black curve shows the fit to the data provided by the model shown in Fig. 2b. As can be seen, the peak of the distribution of error responses is shifted to the right of zero, indicating that subjects had a bias (*M* = 7.8°) in reporting the value of the target stimulus. *(b)* Similar results for all face trials. As can be seen, the peak of the distribution of error responses is again shifted to the right of zero, indicating that subjects had a bias (*M* = 5.0°) in reporting the value of the target stimulus.

### Individual differences in adaptation bias and representation precision are negatively correlated

We then fit the response bias data for each individual subject. We found that adaptation bias did not significantly differ between blocks (repeated measures ANOVA with block and stimulus class as within-subject factors: *F*(1,201) = 0.0160, *p* = 0.9), suggesting that the magnitude of adaptation biases is a stable individual characteristic for the duration of the experiment. For that reason, we again fit the response bias using data from both blocks of each experiment. We then performed a repeated measures analysis of variance to identify the sources of variability in adaptation bias. We observed a small but significant effect of stimulus class (*F*(1, 201) = 34.9, *p* < .001), explaining only 6% of the total variance. The effect of mean subject bias (*F*(201,201) = 1.56, *p* < .001) was more substantial, accounting for 57% of the total variance in bias values. The distribution of individual subject bias was well fit with a Gaussian with a mean of 8.0° for colors (95% CI [0.2°, 18.8°]) and 5.3° for faces (95% CI [−4.9°, 17.1°]). These results are consistent with an individual difference in adaptation bias that is present across face and color stimuli.

We next extracted the width (i.e., precision) of the error distribution for each subject, separately for each block of each experiment. Again, because precision did not significantly differ between blocks (repeated measures ANOVA with block and stimulus class as within-subject factors: *F*(1, 201) = 0.86, *p* = 0.36), we combined data from both blocks of each experiment. We then performed a repeated measures analysis of variance to identify the sources of variability in response precision. We observed a significant effect of stimulus class (*F*(1,201) = 170.1, *p* < .001) accounting for 16.5% of the total variance, with response precision being higher for colors. We also observed a significant effect of mean subject precision (*F*(201,201) = 3.28, *p* < .001) accounting for 64% of the total variance. Furthermore, response precision estimated from *stimulus-match* trials was well correlated with precision estimated from the *adaptation* experiment (color: *r* = 0.45, *p* < 0.001; faces: *r* = 0.44, *p* < 0.001; Fig. 4a,b). This correlation is possibly due to shared sources of variability between experiments. Note, however, that the variance associated with reporting the target on *stimulus-match* trials is smaller than the variance associated with reporting the target in the *adaptation* experiment, possibly due to the fact that the latter is presented briefly and involves a short-term memory component.

**Figure 4:**
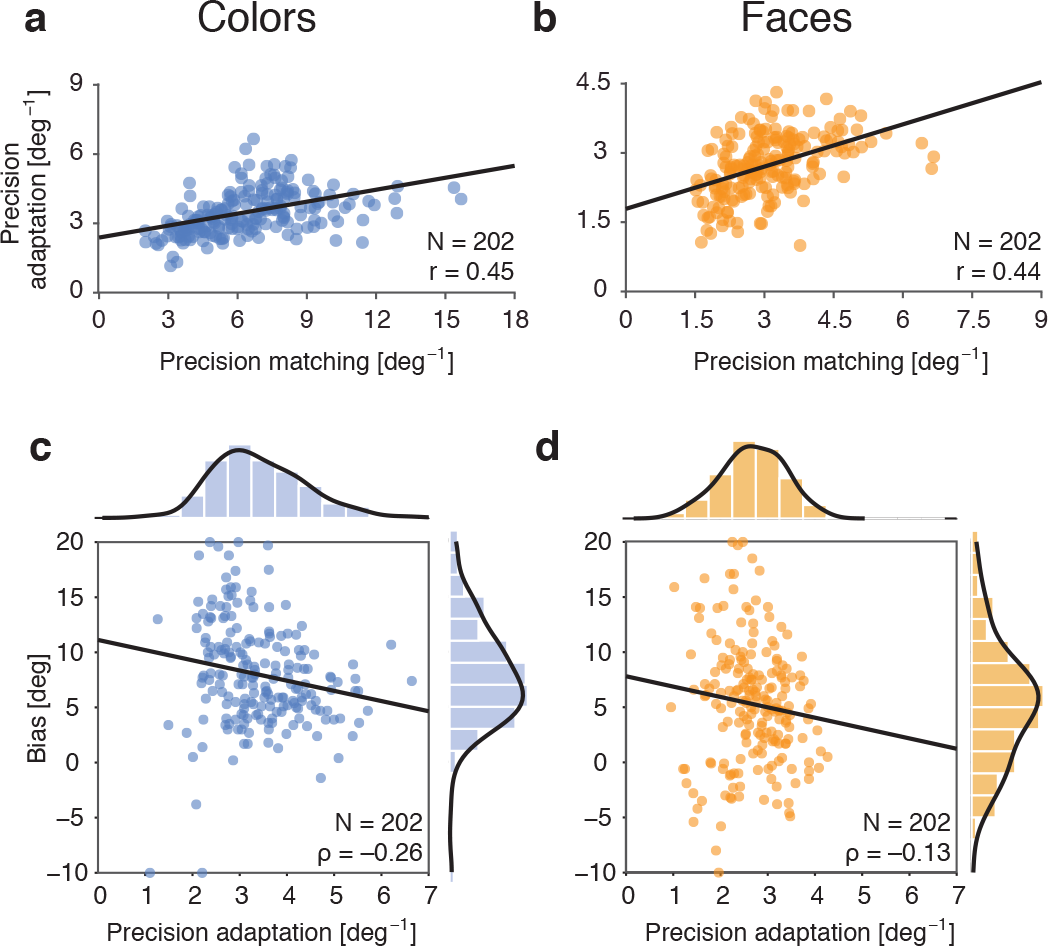
Individual differences in representation precision are negatively correlated with adaptation bias. *(a)* We fit data from each subject collapsed across both blocks of the color stimulus-match experiment and across both blocks of the color adaptation experiment, using the mixture model approach described in Fig. 2. The correlation between representation precision in both experiments was *r* = 0.45. *(b)* Similar to (a), for face stimuli. The correlation between representation precision in in the stimulus-match and adaptation experiments was *r* = 0.44. *(c)* We fit data from each subject collapsed across both blocks of the color adaptation experiment, using the mixture model approach described in Fig. 2. The Spearman’s rank correlation coefficient between representation precision and adaptation bias was *ρ* = −0.26. *(d)* Similar to (c), for face stimuli. The Spearman’s rank correlation coefficient between representation precision and adaptation bias was *ρ* = −0.13.

We then asked if subjects with a lower representation precision are more or less prone to adaptation biases. We investigated the statistical relationship between average representation precision and adaptation bias, both estimated from the *adaptation* experiment. We observed that, across subjects, there was a negative correlation between mean precision and bias (Spearman’s rank correlation, color: *ρ* = −0.26, jackknife resampling 95% CI [−0.28, −0.25], *p* < 0.001; faces: *ρ* = −0.13, jackknife resampling 95% CI [−0.15, −0.12], *p* < 0.001; Fig. 4c,d). This suggests that subjects with lower representation precision exhibit larger biases away from the adapting stimulus, consistent with predictions from a Bayesian observer model.

We also examined the relationship between color bias and face bias across subjects (Fig. 5), finding a weak positive correlation in these scores (Pearson correlation: *r* = 0.22, jackknife resampling 95% CI [−0.21, −0.23], *p* < 0.001). A possible source for this relationship is the shared effect of individual differences in precision. We cannot, however, be certain that this is the entire explanation, as the correlation between face and color bias values remains in partial correlations that attempt to account for individual variation in precision (partial Pearson correlation: *r* = 0.18, jackknife resampling 95% CI [−0.17, −0.19], *p* < 0.001). We considered the possibility that individual differences in reaction time could account for individual differences in adaptation magnitude. The magnitude of perceptual bias decays following cessation of the adaptor (Greenlee et al., 1991). Perhaps subjects who respond quickly tend to experience more bias due to less decay. However, this prediction was not confirmed. For faces, we found no significant relationship between response bias and average response time (Spearman’s rank correlation: *ρ* = 0.07, *p* = 0.30). For colors, the opposite pattern was found: subjects who responded quickly experienced less bias (Spearman’s rank correlation: *ρ* = 0.20, *p* = 0.0047). We do not have a specific mechanism (neural or otherwise) to offer as the basis for this small correlation in induced adaptation bias between materials across subjects.

**Figure 5:**
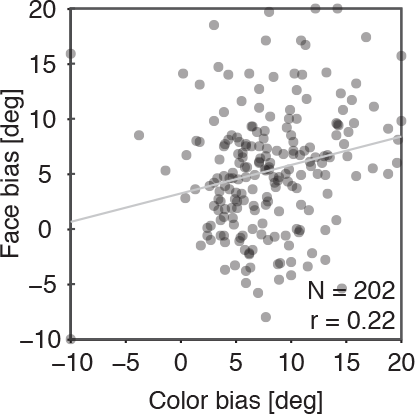
Relationship between color bias and face bias. We fit data from each subject collapsed across both blocks of the color and face adaptation experiments, using the mixture model approach described in Fig. 2. The Pearson correlation coefficient between adaptation bias for colors and faces was *r* = 0.22.

### Replication experiment

We wished to replicate the observed relationship between representation precision and adaptation bias with more trials per subject. An additional group of 472 people were recruited through Amazon Mechanical Turk. From this set, 89 were excluded or abandoned at the color vision test, and 191 during the main experimental trials, leaving 192 subjects for the replication analyses (Tables 3, 4). Subjects performed a slightly modified version of our experiment: two blocks of the *stimulus-match* experiment (30 trials), two blocks of the *delayed-match* experiment (60 trials), and six blocks of the *adaptation* experiment (180 trials). Each subject performed the experiments for only one of the two stimulus classes (faces: 94 subjects; colors: 98 subjects; Fig. 6a). A slightly more saturated version of the stimuli was used in an attempt to increase overall performance (Fig. 6b,c).

**Figure 6:**
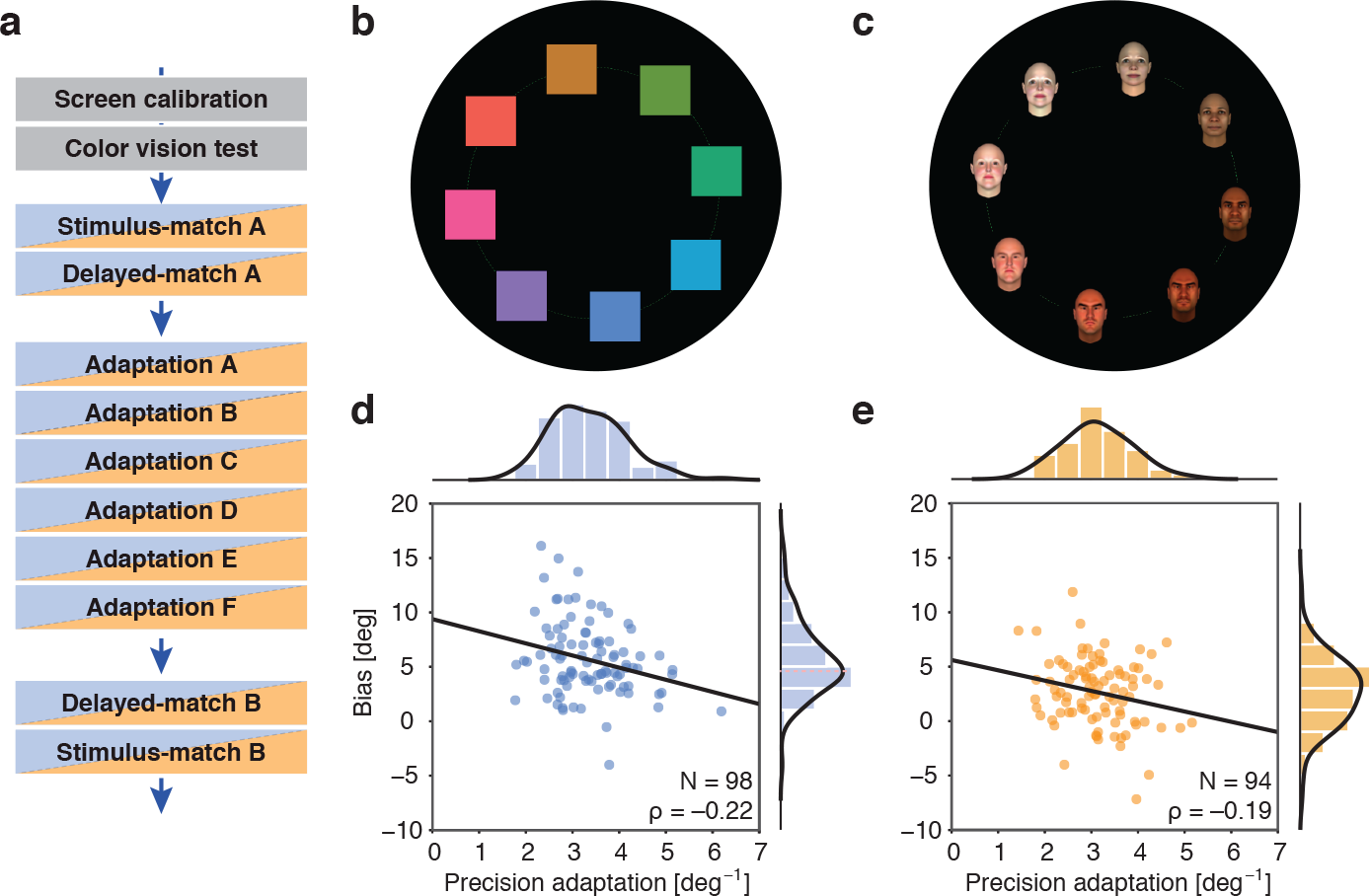
Relationship between representation precision and adaptation bias in the replication experiment. *(a)* Replication experiment structure. Subjects completed a battery of tests designed to estimate representation precision, precision after a delay, and precision and bias after adaptation. Each subject completed two blocks of the stimulus-match and the delayed-match experiments, and six blocks of the adaptation experiment, for either colors or faces. Prior to each experiment, subjects were presented with instructions and a comprehension quiz, and in the first time they performed each block of a given experiment, they also completed 5 practice trials. Each block of the stimulus-match experiment comprised 15 trials. Each block of the delayed-match and adaptation experiments comprised 30 trials. *(b)* Color stimuli. A circular space with 360 stimuli was used. Colors were generated to vary in hue but not in saturation or luminance (HSL space), though saturation was higher than in the first experiment. *(c)* Face stimuli. A circular space with 360 stimuli was used. Faces stimuli varied in age and gender, each forming one axis of the space. To maximize stimulus differences, the extreme points on each axis also varied in identity. *(d)* We fit data from each subject collapsed across all blocks of the color adaptation experiment, using the mixture model approach described in Fig. 2. The Spearman’s rank correlation coefficient between representation precision and adaptation bias was *ρ* = −0.22. *(e)* Similar to (d), for face stimuli. The Spearman’s rank correlation coefficient between representation precision and adaptation bias was *ρ* = −0.19.

After fitting the data using the same approach as described previously (Fig. 2), we analyzed the sources of variability in response precision. Across both color and face experiments, experimental block explained 0. 9% of variance (*F*(5,950) = 3.72, *p* = 0.002) and subject explained 53.5% of variance (*F*(190, 950) = 5.873, *p* < 0.001). We then analyzed the sources of variability in adaptation bias. Similarly, experimental block explained 0.8% (*F*(5,950) = 2.75, *p* = 0.02) and subject explained 47.1% of variance (*F*(190,950) = 4.51, *p* < 0.001). To investigate whether the inverse relationship between representation precision and adaptation biases was replicated, we extracted the bias and precision values for each subject using data from all blocks and calculated the correlation between these two quantities. We again observed that, across subjects, there was a negative correlation between mean precision and bias (color *ρ* = −0.22, jackknife resampling 95% CI [−0.25, −0.20], *p* < 0.001; faces *ρ* = −0.19, jackknife resampling 95% CI [−0.22, −0.17], *p* < 0.001;

## Discussion

We investigated the relationship between individual differences in adaptation bias and response precision for colors and faces. In two cohorts of 202 and 192 subjects recruited through Amazon Mechanical Turk, precision and adaptation bias were found to be stable properties of the observer, with substantial variance in the measurements arising from between-subject differences. Across experiments and materials, we found that lower response precision in an individual was associated with greater perceptual bias.

Two hypotheses have been proposed to reconcile repulsive biases with the Bayesian framework. First, adaptation may induce asymmetries in the likelihood function (Stocker and Simoncelli, 2006a). Combined with a symmetric prior around the adaptor, an asymmetric likelihood can produce estimates that are shifted away from the an observer’s prior (Wei and Stocker, 2015). Alternatively, adaptation may affect the prior itself, according to the expectation that stimulus statistics in recent history match the statistics accumulated over a longer, more distant past (Chopin and Mamassian, 2012). After a prolonged stimulus presentation, a Bayesian observer model would predict biases “away” from this stimulus to maintain balanced statistics in recent history. Note that in the extreme case of infinite variance, the first proposal predicts attraction towards the adaptor (as the likelihood would no longer be asymmetric, estimates would converge again towards the prior, which is centered around the adaptor), while the second continues to predict a bias away from the adaptor and towards the long-term stimulus prior.

A notable finding of our study was the substantial individual variation in the magnitude of perceptual adaptation. This measurement was reproducible across blocks within a testing session. As we did not measure across testing sessions, our measurement likely also contains a component of state variation (although prior studies of individual variation in blur adaptation suggest this component is small; Vera-Diaz et al., 2010). Stable variation across individuals explained almost an order of magnitude more variance in bias measures than did variation within subject across stimulus type (face and color). This indicates a shared mechanism of variation in adaptation magnitude. We find that individual differences in sensory precision provide one such mechanism.

Our results are consistent with a model in which each subject is a Bayesian observer, each of whom differs in the fidelity with which they represent sensory input. We estimated each subject’s precision by measuring response variability. A Bayesian interpretation assumes that response variability in turn reflects (to some degree) individual differences in the precision of sensory encoding. Although quite reproducible, the magnitude of correlation between response precision and adaptation bias was small (accounting for approximately 5% of between-subject variability in adaptation bias). Response precision is therefore an imperfect proxy for sensory precision, or other factors contribute to the substantial between-subject variation in the adaptation effect that is shared across stimulus types.

We recruited subjects through the Amazon Mechanical Turk platform and conducted our experiments using a custom-built website. In addition to allowing a larger sample size, web-based experiments improve subject diversity (Woods et al., 2015). A major challenge of web-based data collection is that subjects may be motivated not to provide a high level of performance, but instead to complete the tasks as quickly as possible to obtain payment. To meet this challenge, our on-line test was designed so that it would be completed most rapidly if the subject produced accurate responses. Additional measures to improve data quality included quizzes to ensure comprehension of the instructions and paying proportionally large bonuses for compliant subjects. While the exclusion of a large proportion of subjects with low accuracy limited the range of precision values we could have measured from our population, we regarded this as an acceptable compromise to exclude subjects who made no attempt to achieve the goals of the measurement.

To conclude, our study provides support for the view that adaptation biases conform with a key property of Bayesian inference: greater perceptual bias should be expected in conditions of lower response precision. While we cannot adjudicate between the conflicting proposals that adaptation affects the prior (Chopin and Mamassian, 2012) or the likelihood (Wei and Stocker, 2015) (or both), our results nonetheless rule out two possibilities. First, that adaptation is not modulated by precision (which would suggest that adaptation bias is not a result of optimal inference). Second, that the perceptual prior is centered on the adaptor when noise is symmetric (which would predict smaller biases in conditions of low precision). Future studies explicitly manipulating expectations independently from sensory evidence, or testing for asymmetries on response distribution, may be conducted to provide further mechanistic evidence.

## Acknowledgements

This research was supported by National Institutes of Health Grants EY021717 (STS and GKA). We thank Alan Stocker for many helpful discussions.

The authors declare no competing commercial interests.

## Author Contributions

Conceptualization, M.G.M., G.K.A., and S.L.T.-S.; Methodology, M.G.M. and G.K.A.; Software and Formal Analysis, M.G.M., M.C., and M.Z.; Visualization, M.G.M. and G.K.A.; Supervision, G.K.A. and S.L.T.-S.; Writing — Original Draft, M.G.M. and G.K.A.; Writing — Review Editing, M.G.M., G.K.A., and S.L.T.-S.; Funding Acquisition, G.K.A. and S.L.T.-S.

## References

Bar, M. (2004). Visual objects in context. Nature Reviews Neuroscience, 5(8):617–629.

Bayes, M. and Price, M. (1763). An essay towards solving a problem in the doctrine of chances. by the late rev. mr. bayes, frs communicated by mr. price, in a letter to john canton, amfrs. Philosophical Transactions (1683-1775), pages 370–418.

Bays, P. M., Catalao, R. F., and Husain, M. (2009). The precision of visual working memory is set by allocation of a shared resource. Journal of vision, 9(10):7–7.

Chopin, A. and Mamassian, P. (2012). Predictive properties of visual adaptation. Current biology, 22(7):622–626.

Clifford, C. W. (2002). Perceptual adaptation: motion parallels orientation. Trends in cognitive sciences, 6(3):136–143.

Fischer, J. and Whitney, D. (2014). Serial dependence in visual perception. Nature neuroscience, 17(5):738.

Fisher, N. I. (1995). Statistical analysis of circular data. Cambridge University Press.

Gibson, J. J. and Radner, M. (1937). Adaptation, after-effect and contrast in the perception of tilted lines. i. quantitative studies. Journal of experimental psychology, 20(5):453.

Greenlee, M. W., Georgeson, M. A., Magnussen, S., and Harris, J. P. (1991). The time course of adaptation to spatial contrast. Vision research, 31(2):223–236.

Inversions, S. (2012). Facegen. Singular Inversions.

Jammalamadaka, S. R. and Sengupta, A. (2001). Topics in circular statistics, volume 5. World Scientific.

Knill, D. C. and Richards, W. (1996). Perception as Bayesian inference. Cambridge University Press.

Lafer-Sousa, R., Hermann, K. L., and Conway, B. R. (2015). Striking individual differences in color perception uncovered by the dress photograph. Current Biology, 25(13):R545–R546.

Levinson, E. and Sekuler, R. (1976). Adaptation alters perceived direction of motion. Vision research, 16(7):779IN7–781.

Portilla, J. and Simoncelli, E. P. (2000). A parametric texture model based on joint statistics of complex wavelet coefficients. International journal of computer vision, 40(1):49–70.

Stocker, A. and Simoncelli, E. P. (2006a). Sensory adaptation within a bayesian framework for perception. Advances in neural information processing systems, 18:1289.

Stocker, A. A. and Simoncelli, E. P. (2006b). Noise characteristics and prior expectations in human visual speed perception. Nature neuroscience, 9(4):578–585.

Summerfield, C. and Egner, T. (2009). Expectation (and attention) in visual cognition. Trends in cognitive sciences, 13(9):403–409.

Vera-Diaz, F. A., Woods, R. L., and Peli, E. (2010). Shape and individual variability of the blur adaptation curve. Vision Research, 50(15):1452–1461.

von Helmholtz, H. (1867). Treatise on physiological optics vol. iii.

Wei, X.-X. and Stocker, A. A. (2015). A bayesian observer model constrained by efficient coding can explain‘anti-bayesian’percepts. Nature neuroscience, 18(10):1509–1517.

Woods, A. T., Velasco, C., Levitan, C. A., Wan, X., and Spence, C. (2015). Conducting perception research over the internet: a tutorial review. PeerJ, 3:e1058.

Zhang, W. and Luck, S. J. (2008). Discrete fixed-resolution representations in visual working memory. Nature, 453(7192):233–235.

